# Expanding the CITE-seq tool-kit: Detection of proteins, transcriptomes, clonotypes and CRISPR perturbations with multiplexing, in a single assay

**DOI:** 10.1101/466466

**Authors:** Eleni Mimitou, Anthony Cheng, Antonino Montalbano, Stephanie Hao, Marlon Stoeckius, Mateusz Legut, Timothy Roush, Alberto Herrera, Efthymia Papalexi, Zhengquing Ouyang, Rahul Satija, Neville E. Sanjana, Sergei B Koralov, Peter Smibert

## Abstract

Rapid technological progress in the recent years has allowed the high-throughput interrogation of different types of biomolecules from single cells. Combining several of these readouts into integrated multi-omic assays is essential to comprehensively understand and model cellular processes. Here, we report the development of <u>E</u>xpanded <u>C</u>RISPR-compatible <u>C</u>ellular <u>I</u>ndexing of <u>T</u>ranscriptomes and <u>E</u>pitopes by <u>seq</u>uencing (ECCITE-seq) for the high-throughput characterization of at least five modalities of information from each single cell: transcriptome, immune receptor clonotypes, surface markers, sample identity and sgRNAs. We demonstrate the use of ECCITE-seq to directly and efficiently capture sgRNA molecules and measure their effects on gene expression and protein levels, opening the possibility of performing high throughput single cell CRISPR screens with multimodal readout using existing libraries and commonly used vectors. Finally, by utilizing the combined phenotyping of clonotype and cell surface markers in immune cells, we apply ECCITE to study a lymphoma sample to discriminate cells and define molecular signatures of malignant cells within a heterogeneous population.

## INTRODUCTION

High-throughput single cell RNA sequencing (scRNA-seq) has rapidly progressed from a tremendous technical achievement to a standard tool for phenotypic interpretation of complex biological systems. scRNA-seq has empowered researchers to deeply phenotype cells, enabling detection of rare cell populations and determination of developmental trajectories of distinct cell lineages. Recently, substantial progress has been made in combining scRNA-seq with genome, epigenome, or perturbation profiling in high throughput assays. Assays that probe the genome and transcriptome of single cells^1,2^ enable detection of single nucleotide variants (SNVs) in matched single-cell genomes and transcriptomes, detection of RNA editing events and detection of expressed, coding mutations. Detection of transcriptomes together with epigenetic modifications including DNA methylation^3-5^ and chromatin accessibility^6^ can link epigenetic regulation of functional elements to gene expression. Additionally, several approaches have recently been reported that allow detection of CRISPR-mediated perturbations along with the transcriptome of single cells, enabling the use of scRNA-seq as an unbiased readout of pooled CRISPR-based genetic screens^7-10^.

scRNA-seq is unbiased in the sense that any polyadenylated transcript can theoretically be captured. In reality, capture of mRNAs is stochastic and inefficient, which reduces the dynamic range of scRNA-seq and gives rise to sparse data. Additionally, for most well studied genes, protein, not mRNA, is the terminal and functional outcome of gene expression, and the detection of mRNA does not always reflect accumulation of the corresponding protein. Previously, we and others have layered detection of proteins on top of scRNA-seq to enable integration of robust and well-characterized protein markers with unbiased transcriptomes of single cells^11,12^. Our method, <u>C</u>ellular <u>I</u>ndexing of <u>T</u>ranscriptomes and <u>E</u>pitopes by <u>seq</u>uencing (CITE-seq) is compatible with oligo-dT based scRNA-seq approaches and enables simultaneous protein detection using DNA oligo-labeled antibodies against cell surface markers. Given that protein levels are typically much higher than corresponding mRNAs, detection of proteins via antibody-derived tags (ADTs) is a more robust measure of gene expression. In a series of experiments, we demonstrated the value of multimodal analysis to reveal phenotypes that could not be discovered by using scRNA-seq alone, as well as the use of CITE-seq for studies of post-transcriptional gene regulation at the single-cell level^11^.

Here, we report the development of <u>E</u>xpanded <u>C</u>RISPR-compatible CITE-seq (ECCITE-seq) for the high throughput characterization of at least five modalities of information from the same single cells. ECCITE-seq adapts CITE-seq^11^ to a 5’ tag-based scRNA-seq assay, integrating clonotype and cell surface marker information to RNA-based cellular phenotypes in immune cells. ECCITE-seq is also compatible with a modified version of Cell Hashing^13^, which allows sample multiplexing, multiplet detection and super-loading of scRNA-seq experiments. Importantly, ECCITE-seq allows the direct detection of single guide RNA (sgRNA) molecules, through a minor modification to the scRNA-seq workflow, which will facilitate high throughput and sensitive single cell perturbation screens compatible with existing guide libraries and commonly used vectors.

## RESULTS

### ECCITE-seq enables the detection of at least five modalities of cellular information from single cells

To enable profiling of protein markers together with V(D)J regions and transcriptomes, we modified our previously described CITE-seq method^11^. Oligos partially complementary to the gel bead-associated template switch oligos in the 10x Genomics 5P / V(D)J kit were covalently conjugated to antibodies as described^13^ and used to label cells. Annealing and extension during the reverse transcription (RT) reaction appends the cell barcode and unique molecular identifier (UMI) to each oligo in parallel with the addition of these sequences to the first strand cDNA copies of cellular mRNAs in the same droplet (Figure 1a) (see methods). Separate detection of differentially expressed proteins, or hashtags, for multiplexing can be accomplished by using different amplification handles. The 10x Genomics 5P workflow requires addition of a soluble poly(dT) oligo to prime RT, opening up the possibility of adding custom RT primers to sequences of interest. sgRNAs have a structure that lends themselves to direct capture: the variable region that guides Cas9 to its target site is at the 5’ end and the 3’ end is an invariant scaffold^14,15^. We leveraged the scaffold as an annealing site for an additional RT primer, which after copying the invariable guide sequence and template switching with the bead-bound oligo acquires a cell barcode and UMI in parallel with other modalities (mRNA, protein tags)(Fig. 1a).

**Figure 1:**
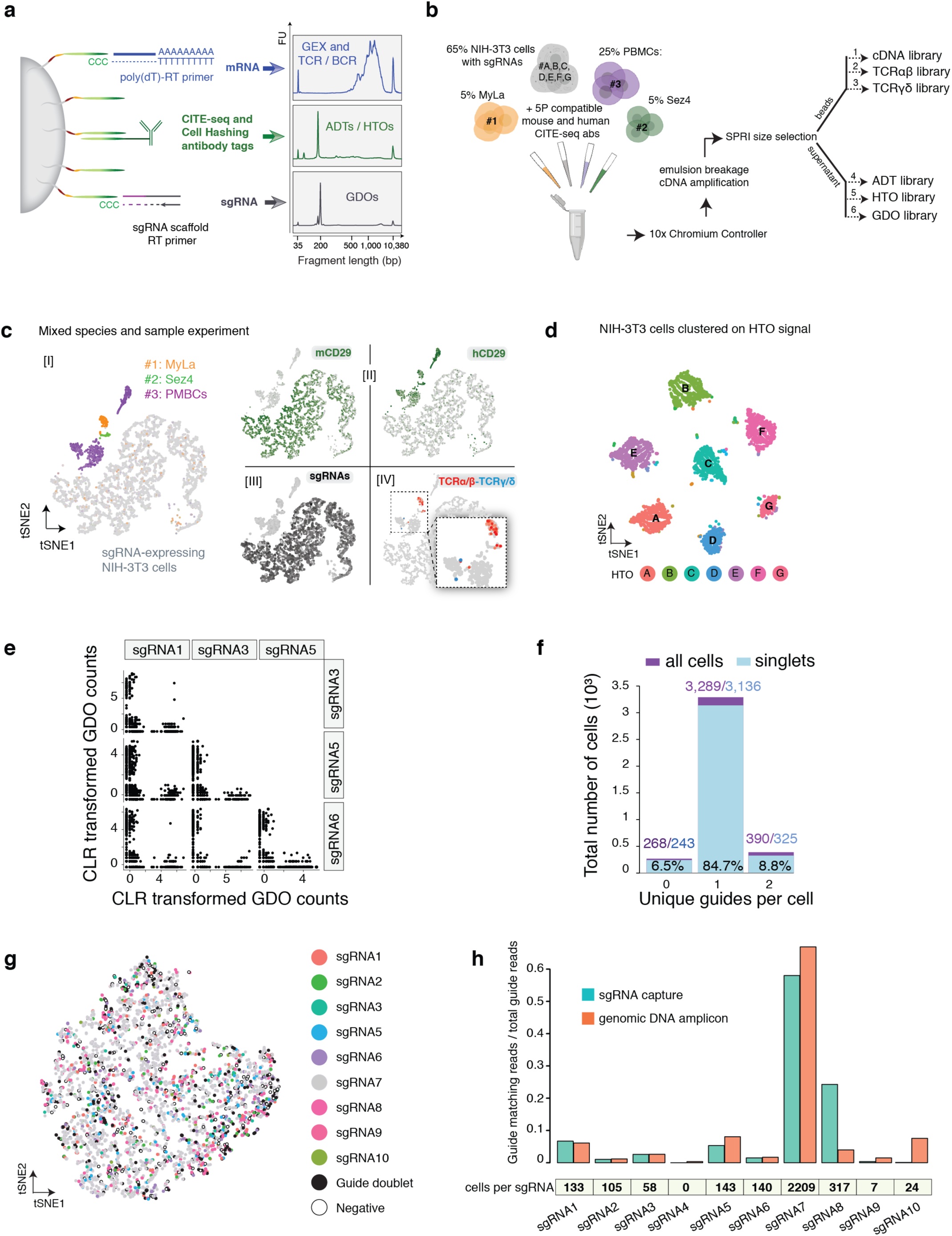
ECCITE-seq allows simultaneous detection of transcriptome, proteins, clonotypes and CRISPR perturbations. a. Schematic overview of the multiple cellular modalities captured by ECCITE-seq. cDNAs derived from mRNA and sgRNA transcripts acquire cell barcodes and unique identifiers through a template switch reaction with the bead-bound oligo, whereas antibody tags harbor sequences that allow direct annealing to the bead oligo. Reverse transcription and amplification yields products with distinct sizes that can be separated and amplified independently (right panel). b. Proof-of-principle ECCITE-seq experiment demonstrating measurement of six cellular modalities. Cells of different origin were stained with CITE-seq and hashing antibodies, washed and combined to a single 10x run. All cells were stained with a mix of anti-human CD29 and anti-mouse CD29 antibodies. NIH-3T3 cells were split into 7 tubes and stained with 7 barcoded hashing antibodies (HTO-A to HTO-G), followed by washing and pooling. MyLa, Sez4 and PBMCs were stained with HTO_1, HTO_2 and HTO_3 respectively. Eventually, a cell mix comprising 65% NIH-3T3 cells, 25% PBMCs, and 5% each MyLa and Sez4 cells was loaded into the 10x Chromium for droplet formation and reverse transcription. After emulsion breakage and cDNA amplification, the distinctly sized products were separated with size selection capture beads and amplified separately. A pool consisting 88% cDNA, 7% GDO, 3% HTO and 2% ADT library was submitted for NGS sequencing. c. Transcriptome-based clustering of single-cell expression profiles of the mixed human and mouse sample, illustrating the 5 modalities of ECCITE-seq: Transcriptome, Cell Hashing [I], Protein [II], sgRNAs [III] and T cell antigen receptors [IV] (most abundant clonotype shown for each cell type, α/β: red, γ/δ: blue). d. t-SNE embedding of the NIH-3T3 hashtag dataset. Cells are colored and labeled based on HTO classification. Seven singlet clusters and 21 cross-sample doublet clusters are present. e. Representative scatterplots of scaled and normalized sgRNA counts of NIH-3T3 cells after removing cell doublets. f. Number of unique sgRNAs detected per cell for all mouse barcodes (purple) or singlets only as defined by cell hashing (blue). g. Transcriptome-based clustering of 3,704 NIH-3T3 singlet cells expressing 10 non-targeting sgRNAs (Supplementary Table 5). Cells were assigned to an sgRNA identity by adapting the HTO demultiplexing model to GDO reads instead of HTO reads. h. sgRNA representation as measured by direct sgRNA capture or genomic DNA amplification of the guide variable region. The histograms represent the ratio of reads matching each sgRNA to the total sgRNA reads and the table inset depicts the cells assigned to each single sgRNA.

To illustrate the detection of six modalities (transcriptome, T cell receptor (TCRα/β and TCRγ/δ), surface protein, sample origin, and sgRNA) in a single experiment, we generated a cell mixture comprising human peripheral blood mononuclear cells (PBMCs), two human T cell lymphoma lines (MyLa and Sez4) and mouse NIH-3T3 cells that had been transduced with a library of nontargeting sgRNA-generating constructs (Fig.1b, Supplementary Table 5). The captured modalities are summarized in Figure 1c. Cell hashtags specific to human cells were used to distinguish the three human samples, and the hashtag distribution was consistent with transcriptome-based clustering (Fig.1c, [I]). CITE-seq antibodies directed against human or mouse CD29 label cells according to their species of origin [II], illustrating the ability of ECCITE-seq to detect differentially expressed proteins within a sample. The addition of an sgRNA scaffold-specific RT primer before emulsion generation (Fig. 1a), and subsequent enrichment PCR (methods) allowed us to capture sgRNA molecules which were present only in mouse cells [III]. Finally, clonotypes for TCRα/β (following 10x protocol) and TCRγ/δ (custom adaptation, see methods) could be distinguished in the PBMC and lymphoma cell clusters [IV]. Prior to encapsulation, the mouse cells in the proof-of-principle experiment were split into seven samples and labeled with separate hashtags. Clustering based on HTO counts revealed the expected major singlet and doublet clusters (Fig. 1d). Consistent with a single cell assay readout, sgRNA counts for different guides show anticorrelation between cell barcodes (representative examples shown in Fig 1e). Importantly, the use of Cell Hashing together with sgRNA detection allowed us to distinguish between apparent “doublets” where cells have been infected with two viruses (n=325), from doublets resulting from co-encapsulation of two cells in the same droplet (n=65) (Fig. 1f). sgRNA capture was highly efficient, with sgRNAs detected in 93.5% of mouse cells (Fig. 1f), and successful over a wide range of sgRNA representation, consistent with proportions of individual guides in the library (Fig.1g,h). We note that this non-targeting library had been maintained in culture for several weeks prior to the experiment, likely allowing population drift. While this is not advised for functional genomics screens, for this proof-of-principle, it served a useful purpose of demonstrating guide capture over a wide range of cellular abundances.

### CRISPR screens with single cell multimodal readout

ECCITE-seq is designed to enable interrogation of single cell transcriptomes together with surface protein markers in the context of CRISPR screens. To illustrate this, we infected HEK-293T cells with a CRISPR library comprising guides targeting genes encoding cell surface markers (CD29 and CD46) and other important signaling molecules (JAK1 and p53), as well as non-targeting controls (Supplementary Table 6). ECCITE-seq on these cells interrogating surface markers, transcriptome and guide RNAs allowed us to determine the effect of introduced sgRNAs on the expression levels of proteins and mRNAs of interest. For CD29, an integrin β subunit highly expressed on the surface of HEK-293T cells, we observed that most cells have detectable levels of protein (Fig. 2a). Protein levels are unchanged in the presence of non-targeting sgRNAs but collapse in cells with detectable levels of sgRNAs directed against ITGB1, the gene encoding CD29. In comparison with CD29 protein, many cells have undetectable levels of ITGB1 mRNA even in the absence of targeting guides, likely reflecting the high drop-out rates of scRNA-seq, and the increased sensitivity that comes with protein detection. mRNA reduction is apparent in cells with targeting sgRNAs but is a less obvious phenotype than observed for protein. Specific loss of gene products was also observed for CD46 protein and JAK1 mRNA, demonstrating the ability to use ECCITE-seq to obtain multimodal readouts for CRISPR screens (Fig 2b). When cells were clustered by sgRNA sequence, reduction of expression within clusters expressing targeting sgRNAs was observed for both mRNA and protein (Fig,2c).

**Figure 2:**
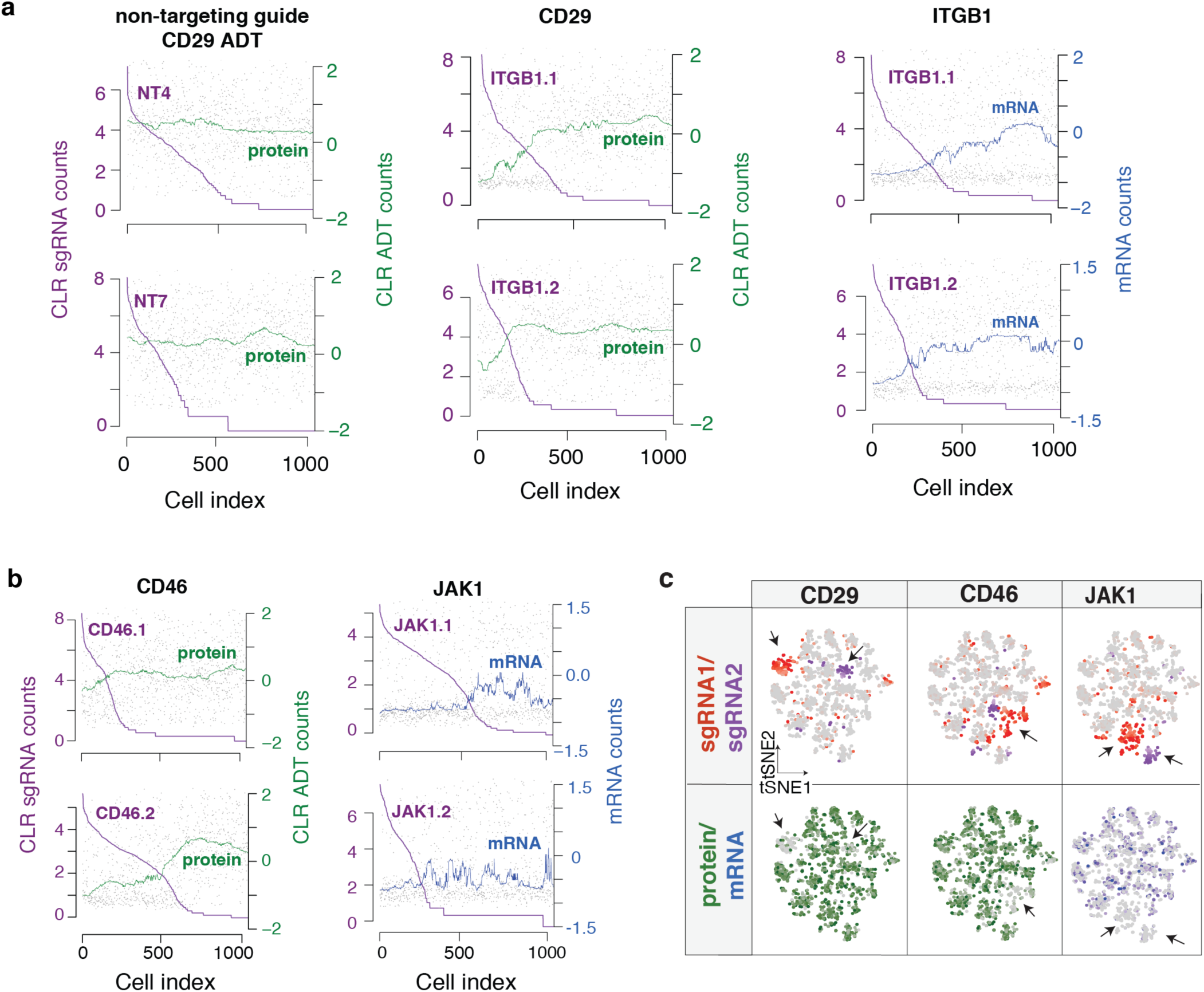
ECCITE-seq experiment demonstrating functional assessment of CRISPR editing in 5,368 HEK cells expressing 10 non-targeting and 11 targeting sgRNAs. a. Levels of target surface marker CD29 or ITGB1 mRNA (moving median smoothing window = 150) in cells expressing targeting and non-targeting sgRNAs, ordered based on decreasing sgRNA counts (purple). Each grey dot represents a cell and where it stands based on sgRNA and protein/mRNA levels, b. Protein or mRNA levels as a function of decreasing sgRNA levels (as described in panel a), using targeting guides against the surface marker CD46 or the intracellular kinase JAK1. c. Cells were clustered based on normalized and scaled sgRNA counts. Highlighted on top are CLR counts for two sgRNAs of each indicated target, and on the bottom counts for respective protein (green) or mRNA (blue).

### ECCITE-seq couples clonotype determination with immunophenotyping

We next constructed a large (49 marker) panel of ECCITE-seq antibodies to deeply profile PBMCs from a healthy donor and a Cutaneous T Cell Lymphoma (CTCL) patient (Fig. 3, Supplementary Fig. 1 and Supplementary Table 1) and prepared libraries for hashtags, ADTs, TCR α/β, TCR γ/δ and transcriptome. After HTO demultiplexing to separate samples and remove doublets, cells were clustered based on transcriptome (Fig. 3a, b and Supplementary Fig. 1). The majority of markers showed enrichment at both RNA and protein level in expected clusters, with a more robust protein signal, consistent with our previous 3’ CITE-seq results^11^. We additionally recovered clonotype information for both the control and CTCL samples and the most abundant clonotypes were located in appropriate T cell clusters. The control sample had 1606 detected clonotypes from 2796 barcodes, with the top CD4+ clonotype (TRB CDR3 sequence= CASSTLQGKETQYF) accounting for 1% of recovered clonotype-associated barcodes. In contrast, in the CTCL sample, a single CD4+ clonotype was dramatically expanded (TRB CDR3 sequence= CSARFLRGGYNEQFF), accounting for 14% of recovered clonotype-associated barcodes (1738 detected clonotypes from 3857 barcodes).

**Figure 3:**
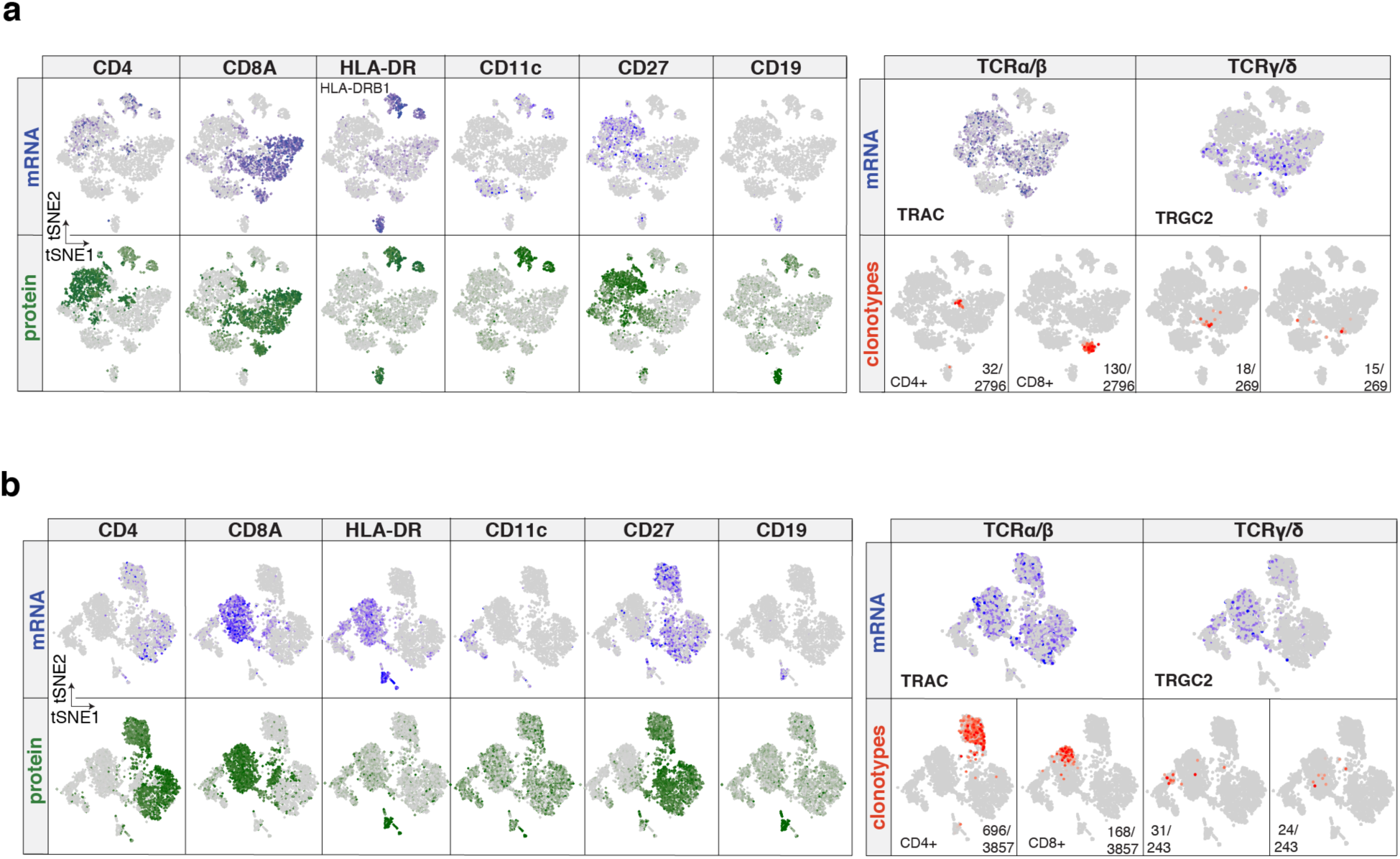
ECCITE-seq on PBMCs from a healthy donor and a CTCL patient (select markers) Transcriptome-based clustering of 4,132 cells from control sample (a) and 4,178 cells from CTCL sample (b) with projected mRNA (blue), corresponding ADT (green) signal for selected CITE-seq antibodies and the two most abundant TCRα/β or TCRγ/δ clonotypes (red).

For further comparative analysis, cells from both samples were computationally merged^16^ and clustering based on either RNA or protein showed agreement in detecting most cell subpopulations and their gene-expression signatures (Fig.4a and Supplementary Fig. 2). TCR sequences were recovered for 76.5% of T cells with the dominant CD4+ clonotypes exhibiting strikingly different clonality, as described above (Fig. 4b). *in silico* gating based on CD3 and CD4 protein levels coupled with clonotypic information enabled differential gene expression analysis comparing malignant cells with polyclonal T cells from both the patient and the healthy donor sample (Fig.4c). This multimodal analysis revealed a distinct gene expression signature of the malignant CTCL cells consistent with prior studies^17^.

**Figure 4:**
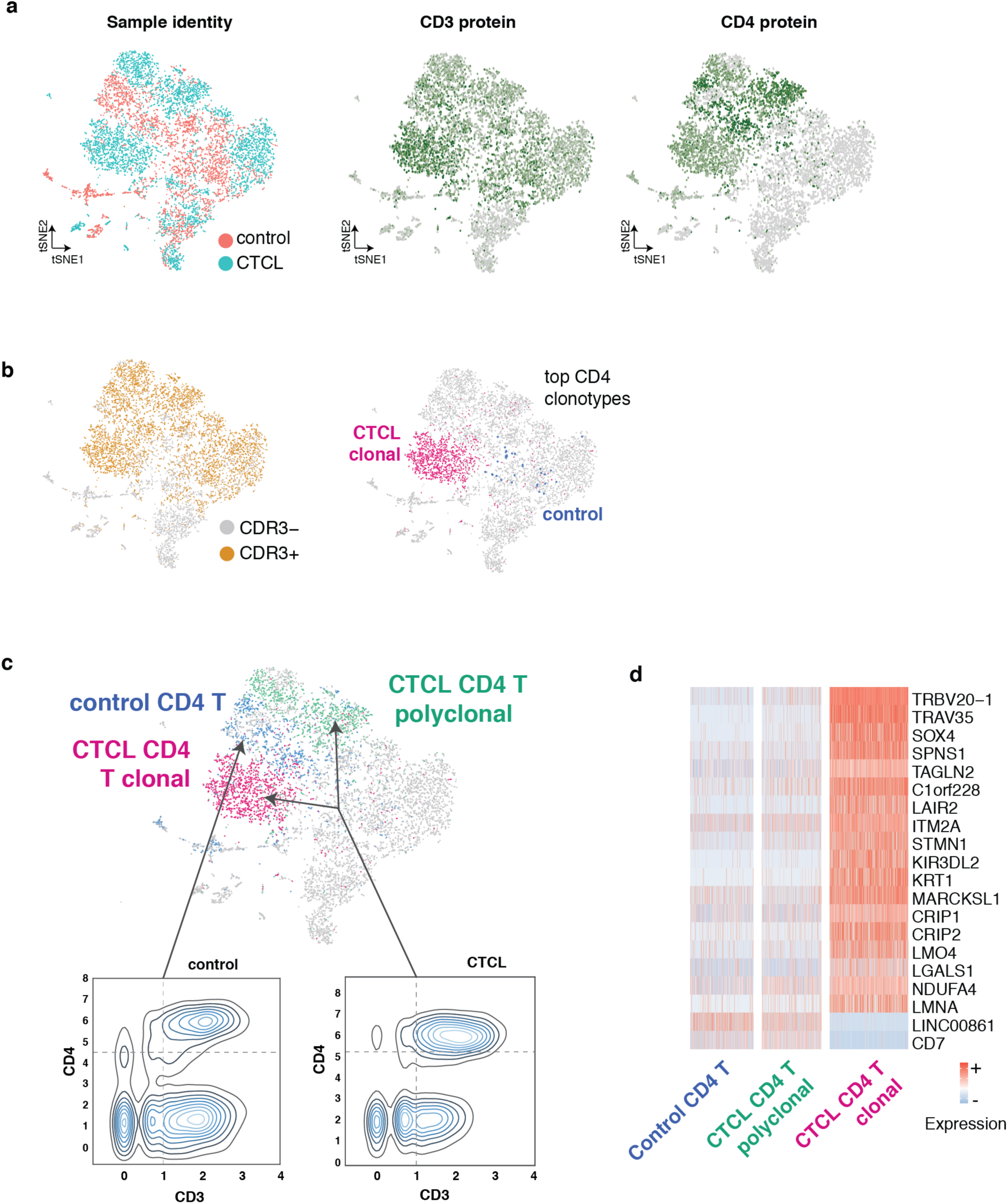
Cell classification based on combined surface marker expression and clonotype determination derived from ECCITE-seq. a. Transcriptome-based clustering of 9,816 cells from both Individuals after removing cell duplicates, merging, depth-normalization, and cell alignment. Highlighted from left to right Is sample Identity, CD3 and CD4 ADT counts, b. Cells with productive TCR rearrangement and most abundant CD4+ TCRα/β clonotype highlighted on the combined dataset, c. In silico gating based on CD3 and CD4 protein counts couple with clonotype determination allows definition of CD4 T cells In control (left) and CTCL (right) samples, d. Heatmap of differentially expressed genes between the CD3+CD4+ defined Th clusters.

## DISCUSSION

The enhancements to the CITE-seq toolkit provided by ECCITE-seq enable detailed phenotypic and functional characterization of single cells. The recovery of clonotype information together with surface protein marker expression allowed fine separation of specific cell populations of interest, enabling careful determination of molecular phenotypes. Analogous to the use of TCR clonotype information in this study, we have recently used expressed mutations to define and further characterize clonal populations in scRNA-seq data-sets (Genotyping of Transcriptomes, GoT^18^), an approach that could readily be combined with ECCITE-seq. The method we describe is inherently customizable and we envisage additional oligo-tagged ligands, such as peptide-loaded MHC complexes for detecting specific TCRs, labeled antigens for detection of antigen specific B cells, or antibodies directed against intracellular proteins being added to future iterations of this system. The combination of Cell Hashing together with direct sgRNA capture will enhance perturbation screens with single cell readouts by allowing the analysis of greater numbers of cells for a given budget by allowing discrimination between single cells with multiple expressed sgRNAs and“doublets”, with 2 cells each with their own guide. The “super-loading” afforded by this knowledge will drive down the per-cell cost of single cell CRISPR screens. The modular nature of ECCITE-seq allows the tailoring of readouts of such screens, potentially allowing the investigator to interrogate panels of transcripts and proteins of interest in response to their perturbations in addition to, or instead of, the transcriptome. Importantly, direct capture of sgRNA molecules overcomes documented problems of barcode swapping events observed with Perturb-seq that have the potential to confound single cell perturbation screens^19-22^. Crop-seq overcomes this issue by incorporating the guide sequence itself into a polyadenylated transcript captured by scRNA-seq, but necessitates the use of non-standard vectors for sgRNA production and does not directly measure the molecules mediating the molecular perturbations^10,19^. Direct capture of sgRNAs has the added benefit of capturing a highly abundant RNA polymerase III transcript, leading to the observed high rates of guide recovery. The use of custom RT primers to specifically capture sgRNAs will allow this approach to be further used for combinatorial screening with different types of guides^23^, and provides a roadmap for targeted capture of additional specific RNA molecules including non-polyadenylated transcripts.

## MATERIALS AND METHODS

### Antibody-oligo conjugates

Antibodies used for CITE-seq and Cell Hashing were obtained as purified, unconjugated reagents from BioLegend (see Supplementary Tables 1, 3 & 4) and were covalently and irreversibly conjugated to barcode oligos by iEDDA-click chemistry as previously described^13^.

### ECCITE-seq on 10x Genomics instrument

Cells (approximately 1.5-2 million per sample) were stained with barcoded antibodies as previously described for CITE-seq and Cell Hashing^11^. Stained and washed cells were loaded into 10x Genomics single cell V(D)J workflow and processed according to manufacturer’s instructions with the following 4 modifications:

1. 12 pmol of an RT-primer complementary to sgRNA scaffold sequences was spiked into the RT reaction (only when sgRNA capture was desired).
2. During the cDNA amplification step, 1 pmol of HTO and GDO additive (the latter only when sgRNA capture was desired) and 2 pmol of ADT additive were spiked into the cDNA amplification PCR.
3. Following PCR, 0.6X SPRI was used to separate the large cDNA fraction derived from cellular mRNAs (retained on beads) from the ADT-, HTO- and GDO-containing fraction (in supernatant). The cDNA fraction was processed according to the 10x Genomics Single Cell V(D)J protocol to generate the transcriptome library and the TCRα/β library. To amplify TCRγ/δ transcripts we implemented a similar to TCRα/β approach with a two-step PCR: during target enrichment 1 we used SI-PCR (AATGATACGGCGACCACCGAGATCTACACTCTTTCCCTACACGACGCTC) and a mix of R1_hTRDC (AGCTTGACAGCATTGTACTTCC) and R1_hTRGC (TGTGTCGTTAGTCTTCATGGTGTTCC), followed by target enrichment 2 with a generic P5 oligo (AATGATACGGCGACCACCGAGATCTACAC) and a mix of R2_hTRDC (TCCTTCACCAGACAAGCGAC) and R2_hTRGC (GATCCCAGAATCGTGTTGCTC). cDNA and TCR (α/β and γ/δ) enriched libraries were further processed according to the 10x Genomics Single Cell V(D)J protocol.
4. An additional 1.4X reaction volume of SPRI beads was added to the ADT/HTO/GDO fraction from step 3, to bring the ratio up to 2.0X. Beads were washed with 80% ethanol, eluted in water, and an additional round of 2.0X SPRI performed to remove excess single stranded oligonucleotides carried over from the cDNA amplification reaction. After final elution, separate PCR reactions were set up to generate the ADT library (SI-PCR and RPI-x primers), the hashtag library (SI-PCR and D7xx_s) and the GDO library (SI-PCR and Next_nst_x). The ADT and HTO libraries were prepared as previously described^13^. Following the cDNA amplification, the sgRNA sequences are converted to an Illumina library by amplification with smRNA_nst_x (v3) or Next_nst_x (v4) together with the SI-PCR primer. Prior to the final library PCR, sgRNA molecules can be further enriched by performing extra rounds of amplification with GDO additive and SI-PCR primers.

A pool consisting 88% cDNA, 7% GDO, 3% HTO and 2% ADT library, was sequenced in an Illumina HiSeq 2500 using a 2×25 recipe Rapid Run. A detailed and regularly updated point-by-point protocol for CITE-seq, Cell Hashing, ECCITE-seq and future updates can be found at www.cite-seq.com

### Cells

Patient and control samples were collected at New York University Langone Medical Center in accordance with protocols approved by the New York University School of Medicine Institutional Review Board and Bellevue Facility Research Review Committee (IRB#i15-01162). CTCL patients were diagnosed according to the WHO classification criteria. After written informed consent was obtained, peripheral blood samples were harvested. Peripheral blood mononuclear cells (PBMCs) were isolated from the blood of patients and healthy controls by gradient centrifugation using Ficoll-Paque^™^ PLUS (GE Healthcare) and Sepmate^™^-50 tubes (Stemcell). Buffy coat PBMCs were collected and washed twice with PBS 2% FBS and cryopreserved in freezing medium (40% RPMI 1640, 50% FBS and 10% DMSO). Cryopreserved PBMCs were thawed for 1-2 minutes in a 37° C water bath, washed twice in warm PBS 2% FBS and resuspended in complete medium (RPMI 1640 supplemented with 10% FBS and 2mM L-Glut). Control and CTCL PBMCs were stained with a 49 antibody panel (Supplementary Table 1) and Cell Hashing antibodies (Supplementary Table 4), before loading into two separate 10x lanes. The Sez4 cell line is derived from the blood of an SS patient^24^, and the MyLa 2059 line is derived from a plaque biopsy sample of an MF patient^25^. Sez4 cells were cultured in RPMI 1640 medium with 2mM L-glutamine, 1% Pen/Strep, 500 units/ml of rh IL-2 (Corning), and 10% human serum. MyLa 2059 cells were cultured in RPMI 1640 medium with 2mM L-glutamine, 1% Pen/Strep, and 10% fetal bovine serum. All cells were incubated at 37°C, 5% CO_2_ in a humidified incubator. The cells were cryopreserved in 90% FBS 10% DMSO and aliquots of 1-1.5 million cells were thawed on the day of the experiment. Peripheral blood mononuclear cells (PBMCs) were obtained cryopreserved from AllCells (USA) and used immediately after thawing. NIH-3T3 (mouse) cells expressing non-targeting sgRNAs were maintained according to standard procedures in Dulbecco’s Modified Eagle’s Medium (Thermo Fisher, USA) supplemented with 10% fetal bovine serum (Thermo Fisher, USA) and 1*μ*g/ml puromycin, at 37°C with 5% CO_2_.

#### Lentivirus production and transduction

The sgRNAs were individually synthesized (Integrated DNA Technologies) and cloned into the lentiviral transfer vector LentiCRISPR v2^26^ (Addgene Plasmid: 52961). Equal amounts of each sgRNA vector were mixed and packaged into lentiviral particles through transfection with packaging plasmids in HEK293FT cells, as previously described^27^. For transduction of NIH-3T3, the lentiviral guide pool consisted of 10 non-targeting mouse guides (Supplementary Table 5); for transduction of HEK293FT, the guide pool consisted of 10 non-targeting and 11 gene-targeting human guides (Supplementary Table 6). NIH-3T3 and HEK293FT cells were infected at MOI = 0.05 and selected and maintained in 1*μ*g/ml puromycin. NIH-3T3 cells used in the proof-of-principle experiment were maintained in culture for several weeks, allowing drift in the representation of guides.

### Single-cell data processing

Fastq files from the 10x libraries with four distinct barcodes were pooled together and processed using the cellranger count pipeline. Reads were aligned to the GRCh38 (human healthy and CTCL PBMC datasets) or hg19-mm10 concatenated reference (human-mouse experiment). For ADT, HTO and GDO quantification, we used a previously developed tag quantification pipeline, available at https://github.com/Hoohm/CITE-seq-Count, run with default parameters (maximum Hamming distance of 1). For the TCR libraries, fastq files from the 10x libraries with four distinct barcodes were pooled together, processed using the cellranger vdj pipeline and reads were aligned to the GRCh38 reference genome.

#### Seurat

Normalization and downstream analysis of RNA data were performed using the Seurat R package (version 2.3.0, Satija Lab) which enables the integrated processing of multi-modal single cell datasets. ADT, HTO and GDO raw counts were normalized using centered log ratio (CLR) transformation, where counts were divided by the geometric mean of the corresponding tag across cells, and log-transformed^11^. For demultiplexing based on HTO or GDO counts we used the HTODemux function within the Seurat package as described^13^, with the default percent cutoff = 0.999. For the TCR libraries, productive clonotypes were filtered and their raw counts were inserted into the Seurat object under a new assay slot. Raw counts were normalized using centered log ratio (CLR) transformation and scaled. For comparison between the healthy donor and CTCL data, both Seurat objects were merged and depth-normalized when performing cell alignment (or batch normalization) using RunCCA with a default parameter of 30 canonical vectors^16^. The top 10 aligned components were used for visualization with t-SNE as well as clustering with modularity optimization. The top 20 genes upregulated in each cluster (FindAllMarkers) was used to label the cluster. For ADT clustering, distance matrices of the combined object were computed before generating t-SNE plots.

#### Definition of CD4 T cells and Malignant clone

In analogous strategy to what is used for data visualization in flow cytometry, biaxial KDE plots were made using log(ADT+1) of CD3 and CD4. Cells in both samples were gated at a threshold >=4.5 (log scale) for CD4 ADT and >=1.0 (log scale) for CD3 ADT, defining CD4+ T cells. CTCL Malignant cells were defined as CD4 T cells that possessed the most abundant TCRβ CDR3 amino acid sequence, CSARFLRGGYNEQFF, while CTCL CD4 polyclonal cells were CD4 T cells that did not possess this sequence.

#### Single-cell differential analysis

Comparisons were done using Wilcoxon rank sum test (FindMarkers) between “CTCL Malignant” and “CTCL CD4 polyclonal” as well as between “CTCL Malignant” and “control CD4 Normal”. Significant genes were defined using q-value < 0.05 and |avg_log_2_FC| > 1.0. All Ribosomal Protein (^^^RP[SL][:digit:]) genes as well as Y, X-escapee and X-variable genes were removed from the differentially expressed list. Heatmaps were made using the union of both sets of significant genes.

## ACKNOWLEDGEMENTS

We thank Drs. Odum and Kaltoft for the kind gift of cell lines. Patient samples were obtained with the help of Michal Bar Natan Zommer and Jo-Ann Latkowski. We thank Lu Yang for helpful discussions.

---

**Supplementary Figure 1:**
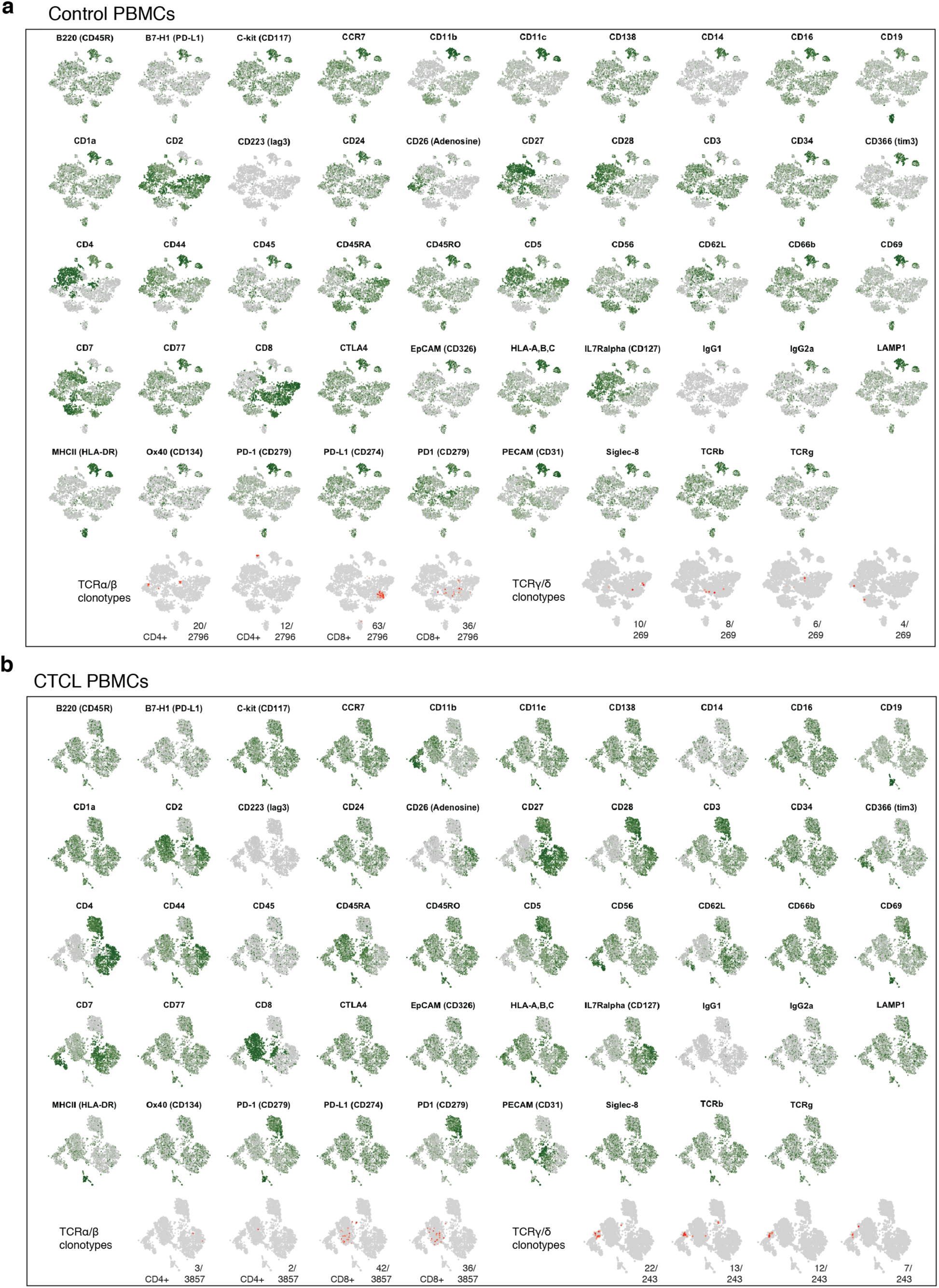
ECCITE-seq on PBMCs from a healthy donor and a CTCL patient (full antibody panel). Normalized and scaled ADT counts of the 49 antibody panel and additional TCRα/β or TCRγ/δ clonotypes (red), projected on the transcrlptome-based t-SNE plots from Fig. 3b and c.

**Supplementary Figure 2:**
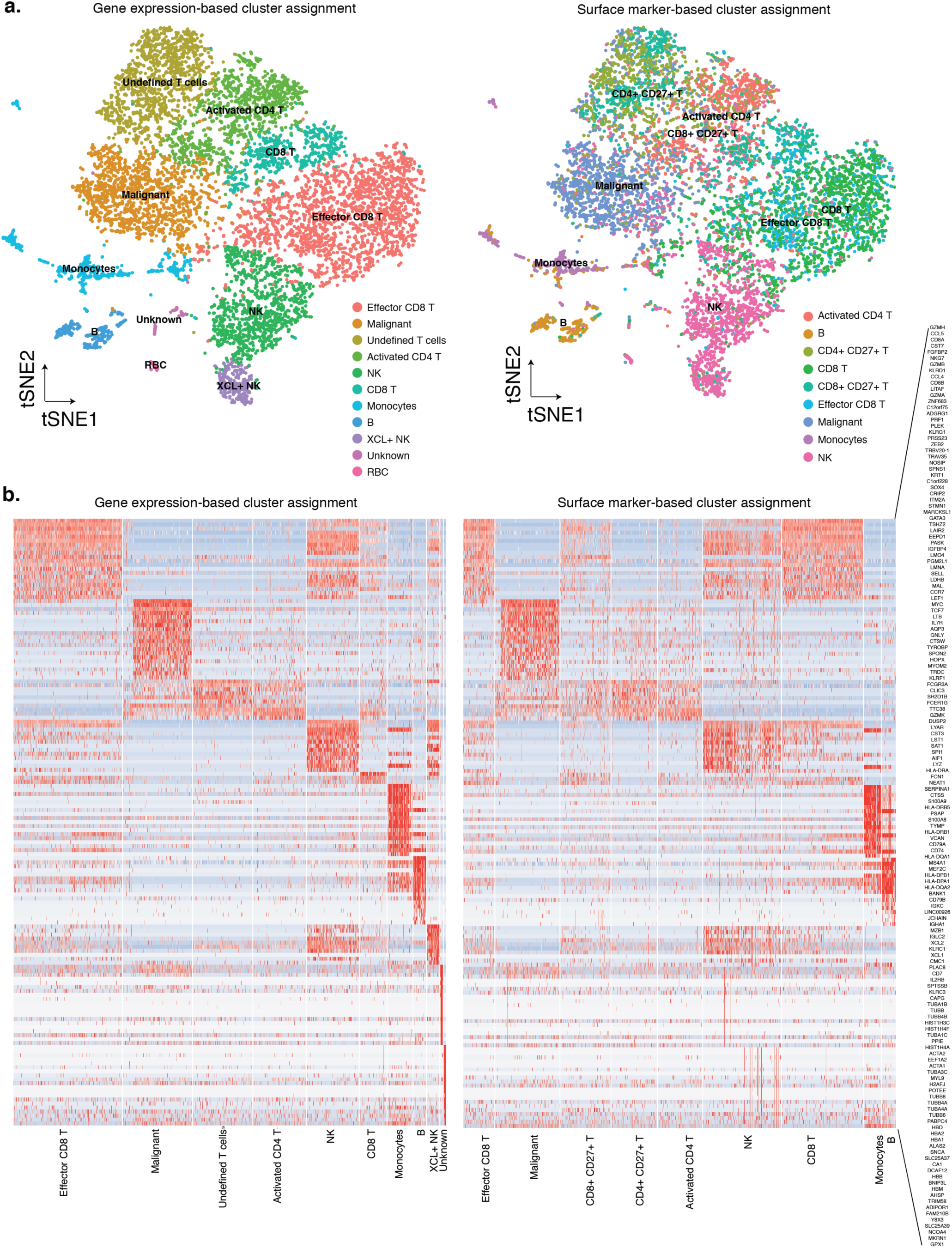
Gene expression and surface marker-based clustering of the combined dataset. a. Unsupervised clustering of PBMCs from both healthy and CTCL donors (n=9,816) after removing cell duplicates, merging, depth-normalization, and cell alignment. Each cell was colored and labelled based on unsupervised clustering information on gene expression (left) or surface marker (right), b. Heatmap of genes differentially expressed across gene expression-based (left) or surface marker-based (right) cluster assignments.

**Supplementary Table 1:**
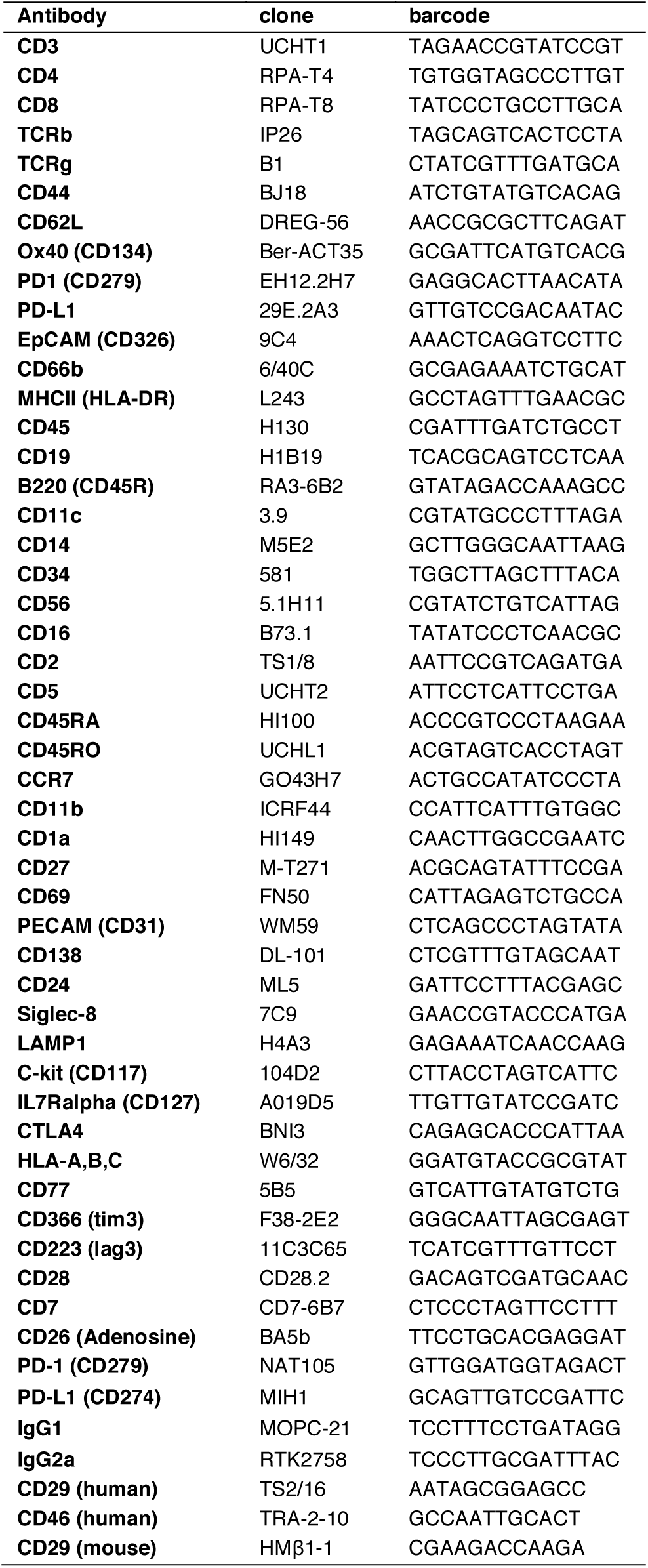
CITE-seq antibody-oligo conjugates

**Supplementary Table 2:**
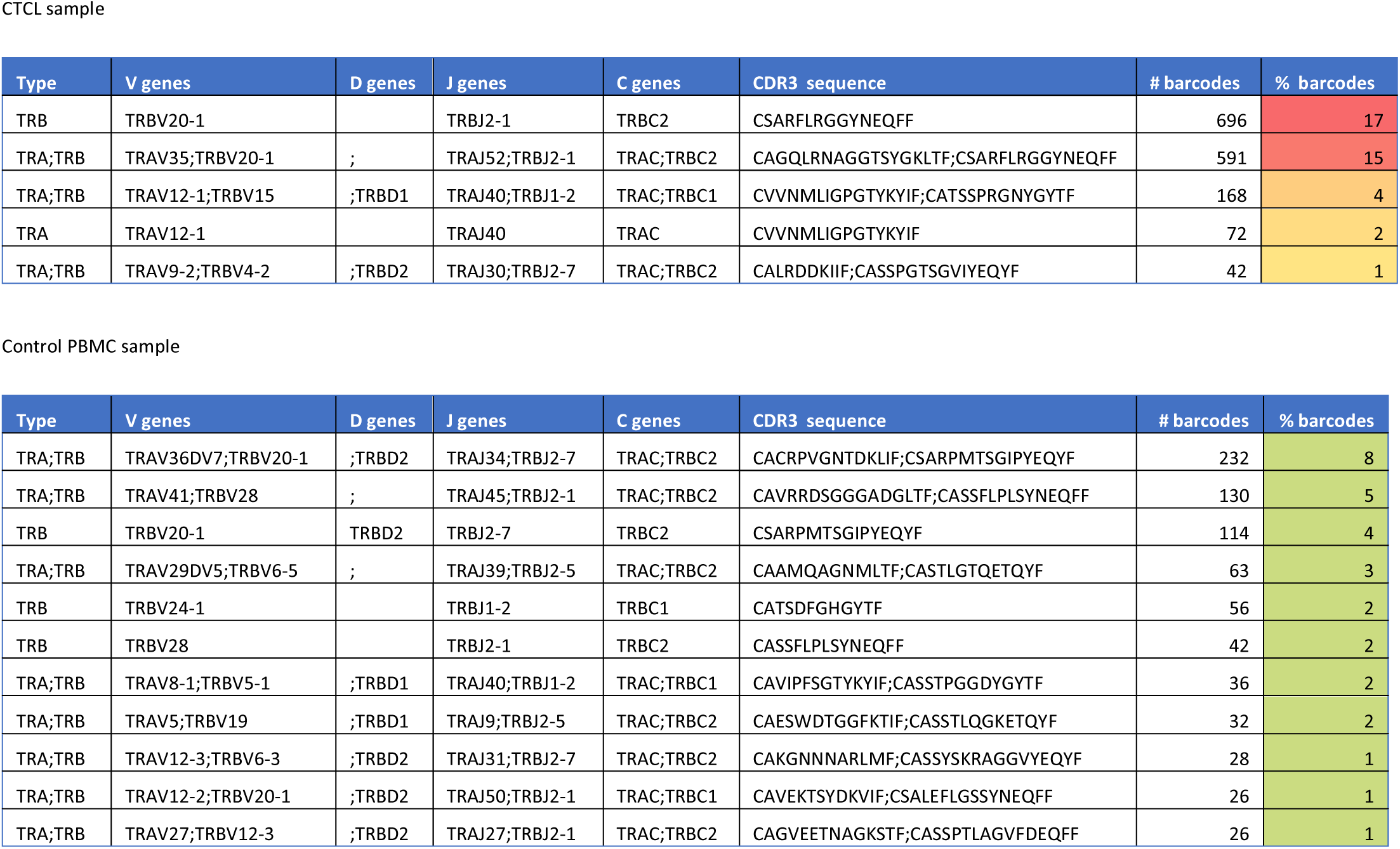
Top clonotypes detected

**Supplementary Table 3:** Mouse hashing antibody-oligo conjugates

| ANTIBODY COCKTAIL | HASHTAG | BARCODE |
| --- | --- | --- |
| CD29 (HMB1-1)<br>CD45 (30-F11)<br>B2M (A16041A) | HTO_A | AGGACCATCCAA |
|  | HTO_B | TCGATAATGCGA |
|  | HTO_C | GAGGCTGAGCTA |
|  | HTO_D | GTGTGACGTATT |
|  | HTO_E | ACTGTCTAACGG |
|  | HTO_F | CACATAATGACG |
|  | HTO_G | TAACGACGTGGT |

**Supplementary Table 4:** Human hashing antibody-oligo conjugates

| ANTIBODY COCKTAIL | HASHTAG | BARCODE |
| --- | --- | --- |
| CD298 (LNH-94)<br>B2M (2M2) | HTO_1 | ACATGTTACCGT |
|  | HTO_2 | AGCTTACTATCC |
|  | HTO_3 | TATCACATCGGT |
|  | HTO_4 | ACTGTCTAACGG |
|  | HTO_5 | CACATAATGACG |
|  | HTO_6 | TAACGACGTGGT |

**Supplementary Table 5:**
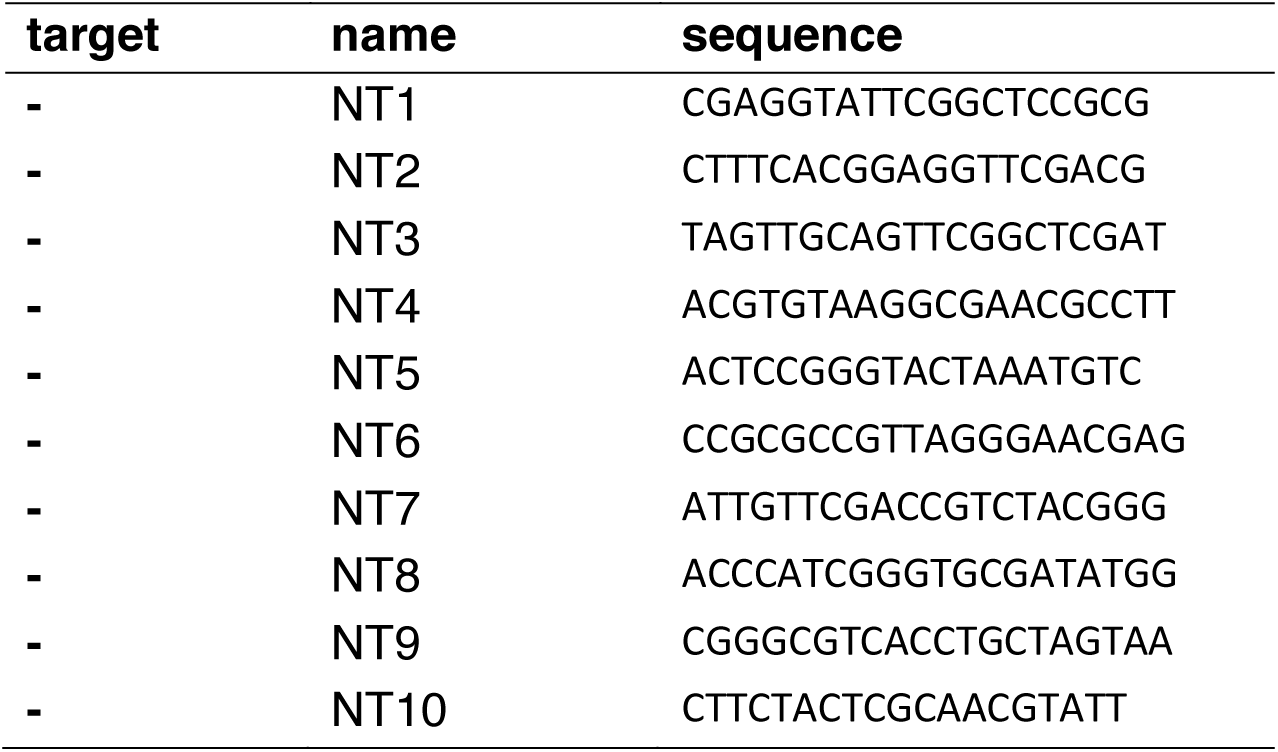
Mouse guide sequences

**Supplementary Table 6:**
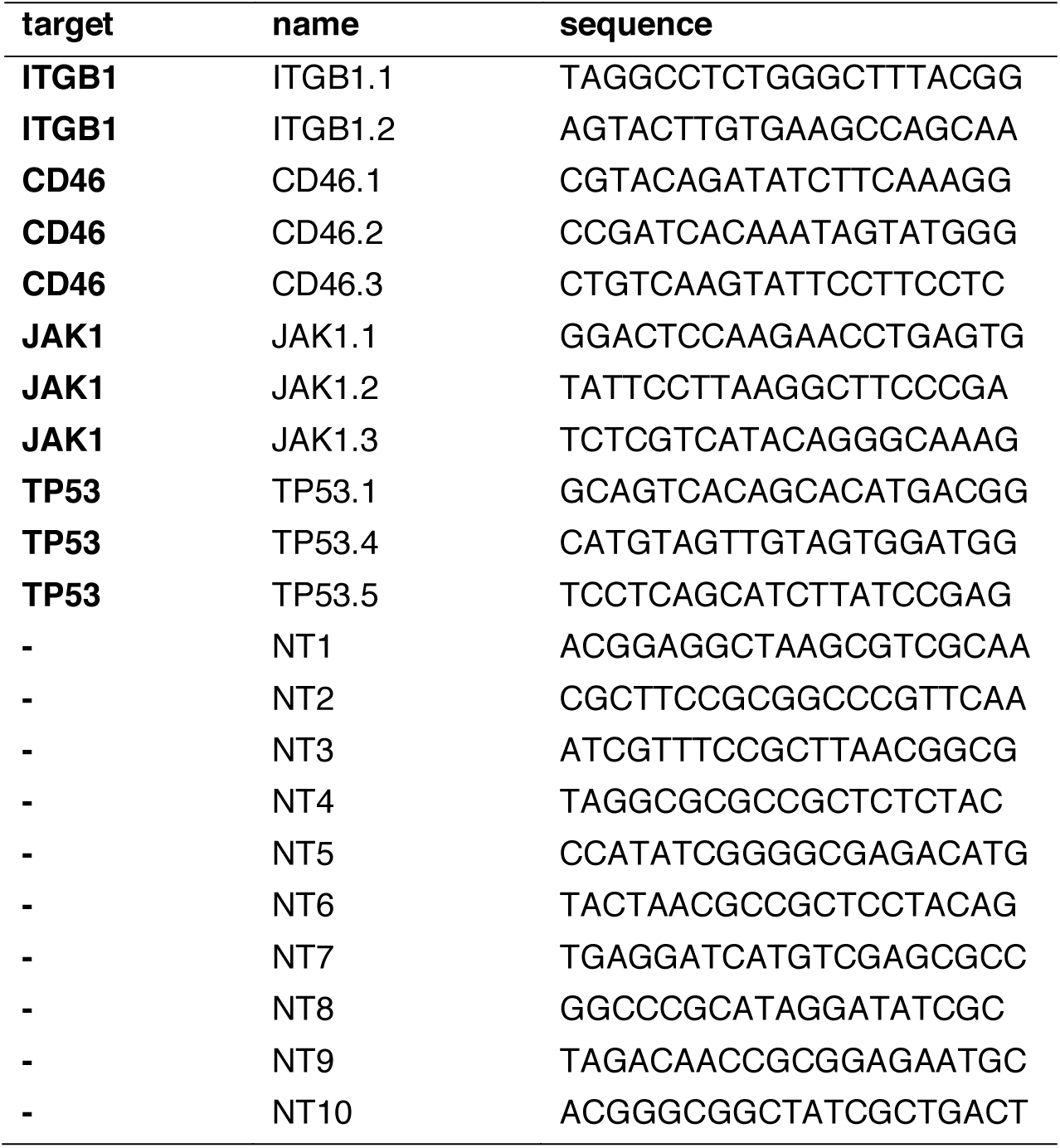
Human guide sequences

